# Monitoring norepinephrine release *in vivo* using next-generation GRAB_NE_ sensors

**DOI:** 10.1101/2023.06.22.546075

**Authors:** Jiesi Feng, Hui Dong, Julieta Lischinsky, Jingheng Zhou, Fei Deng, Chaowei Zhuang, Xiaolei Miao, Huan Wang, Hao Xie, Guohong Cui, Dayu Lin, Yulong Li

## Abstract

Norepinephrine (NE) is an essential biogenic monoamine neurotransmitter, yet researches using prototype NE sensors were limited by their low sensitivities. Here, we developed next-generation versions of GPCR activation-based NE sensors (GRAB_NE2m_ and GRAB_NE2h_) with a superior response, high sensitivity and selectivity to NE both *in vitro* and *in vivo*. Notably, these sensors can detect NE release triggered by either optogenetic or behavioral stimuli in freely moving mice, producing robust signals in the locus coeruleus and hypothalamus. With the development of a novel transgenic mouse line, we recorded both NE release and calcium dynamics with dual-color fiber photometry throughout the sleep-wake cycle; moreover, dual-color mesoscopic imaging revealed cell type‒specific spatiotemporal dynamics of NE and calcium during sensory processing and locomotion. Thus, these new GRAB_NE_ sensors are valuable tools for monitoring the precise spatiotemporal release of NE *in vivo*, providing new insights into the physiological and pathophysiological roles of NE.

## Introduction

Norepinephrine (NE) is a monoamine neurotransmitter that plays essential roles in both the central and peripheral nervous systems, including regulating the sleep-wake cycle^1^, the stress response^2^, attention^3^, sensory processing^4^, heart rate^5^, and blood pressure^6^. Previous methods for measuring NE release *in vivo* relied on either specific—but slow— microdialysis coupled with biomedical identification^7–12^ or rapid—but less specific— electrochemical methods^13–16^ such as fast-scan cyclic voltammetry. The development of CNiFER sensors^17^ and FRET-based sensors^18–20^ provided a means to optically measure NE release with high specificity and temporal resolution; however, the use of these tools has been limited by their undesirable immunogenicity, relatively poor cell-type specificity, and/or narrow dynamic range.

We previously developed a set of genetically encoded G protein‒coupled receptor (GPCR) Activation‒Based (GRAB) NE sensors called GRAB_NE1m_ and GRAB_NE1h_ in which the NE-induced conformational change in the α2AR noradrenergic receptor drives a fluorescence change in circular permutated EGFP (cpEGFP)^21^. These fluorescent sensors outperformed the above traditional methods in sensitivity, selectivity, spatiotemporal resolution, and non-invasiveness. However, the first generation of GRAB_NE_ sensors, GRAB_NE1m_ and GRAB_NE1h_, still had limitations on either molecular sensitivity or selectivity for NE. To further improve these sensors, we developed next-generation GRAB_NE2m_ and GRAB_NE2h_ sensors with 4- fold maximum fluorescence response, nanomolar affinity and more than 200-fold distinguish ability to dopamine (DA). Importantly, these new sensors have rapid kinetics and negligible downstream coupling; in addition, when expressed *in vivo* they produce an up-to 5-fold stronger signal in response to optogenetically and behaviorally stimulated NE release compared to the previous GRAB_NE_ sensors. Moreover, we generated a Cre-dependent transgenic mouse line expressing both green fluorescent GRAB_NE2m_ and the red fluorescent calcium indicator jRGECO1a^22^, which we then used to simultaneously monitor cell type‒specific NE release and calcium dynamics during the sleep-wake cycle, sensory processing, and locomotion. Together, these robust new tools can be used to measure noradrenergic activity under a wide range of physiological and pathophysiological conditions, providing important new insights into the functional role of NE in both health and disease.

## Results

### Optimization and in vitro characterization of next-generation GRAB_NE_ sensors

Our previous fluorescent NE sensors GRAB_NE1m_ and GRAB_NE1h_ reported endogenous NE release with high spatiotemporal resolution^21^; however, when used *in vivo* these sensors have a relatively modest change in fluorescence (∼ 5% in response to optogenetic stimulation in the locus coeruleus), possibly due to low NE sensitivity. GRAB sensors respond to ligand binding by transducing the receptor’s conformational change into a change in cpEGFP fluorescence. To increase the sensitivity of our GRAB_NE_ sensors, we systematically performed site-directed mutagenesis of more than 20 amino acids in GPCR backbone and cpEGFP of GRAB_NE1h_ and then screened the fluorescence responses of more than 400 candidate sensors in HEK293T cells using a high-content imaging system. Among these candidates, one sensor, which we call GRAB_NE2m_, produced the highest change in fluorescence (ΔF/F_0_) in response to NE (Figure 1A). To further increase the sensor’s affinity, we screened sites related to ligand binding and G protein coupling and identified a high-affinity sensor, which we call GRAB_NE2h_ (Figure 1A).

**Figure 1.**
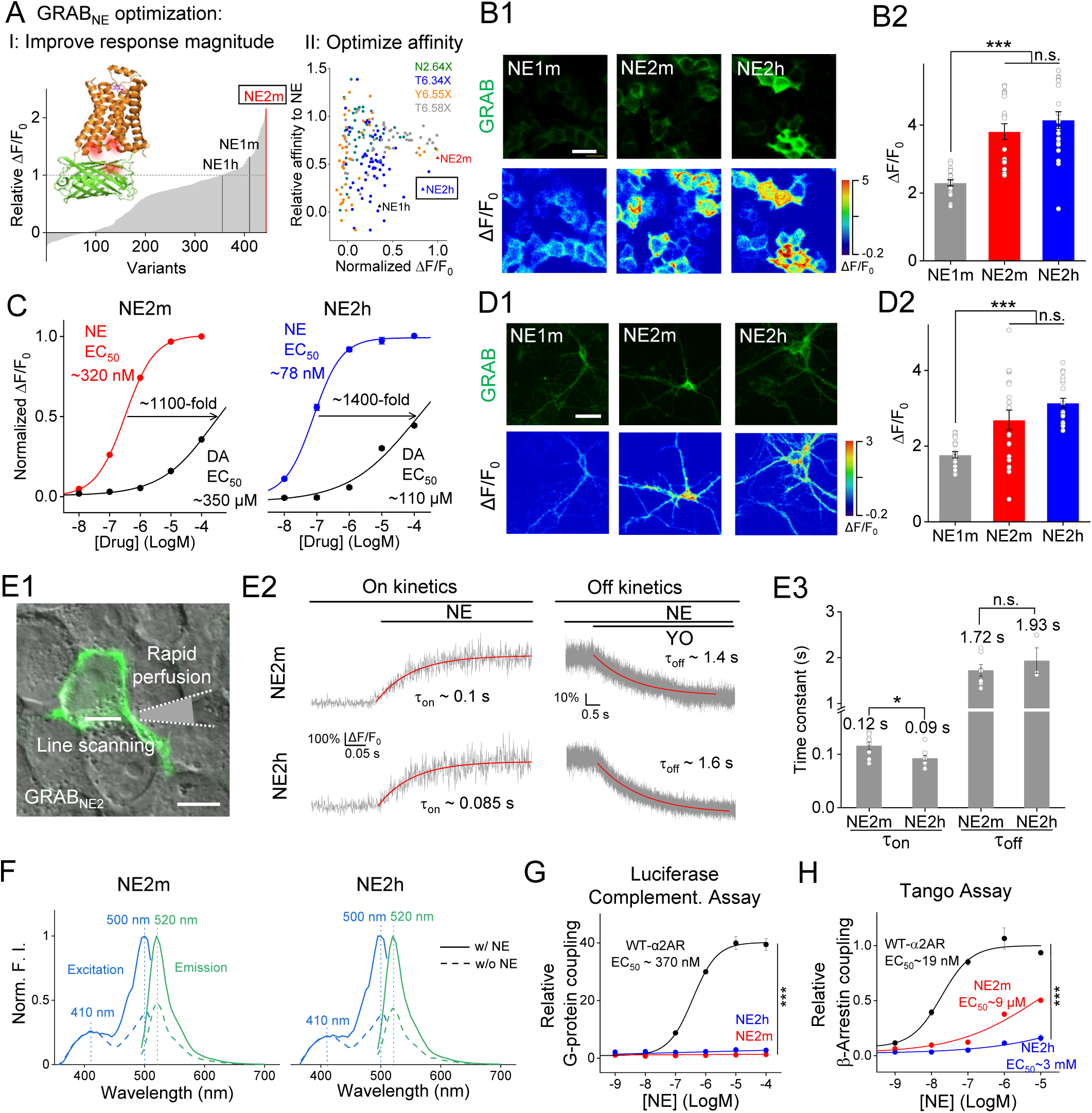
Optimization and *in vitro* characterization of next-generation GRAB_NE_ sensors. (A) I: Optimization of GRAB_NE_ sensors by introducing random mutations at the interface between α2AR and cpEGFP. The first-generation GRAB_NE1m_ and GRAB_NE1h_ sensors, as well as the next-generation GRAB_NE2m_ sensor, are indicated. II: Further optimization to yield GRAB_NE_ sensors with increased ligand affinity, with relative ligand affinity plotted against ΔF/F_0_ (normalized to GRAB_NE2h_). The various mutations are indicated, as well as GRAB_NE1h_, GRAB_NE2m_, and GRAB_NE2h_ sensors. (B1) Images of cultured HEK293T cells expressing GRAB_NE1m_, GRAB_NE2m_, or GRAB_NE2h_. The top row shows baseline fluorescence, while the bottom row shows the change in fluorescence (ΔF/F_0_) in response to 100 μM NE. (B2) Summary of ΔF/F_0_; n = 20 cells from 3 cultures per group. (C) Normalized dose-response curves for GRAB_NE2m_ (left) and GRAB_NE2h_ (right) in response to NE and DA, respectively, in cultured cortical neurons. The corresponding EC_50_ values and fold change in EC_50_ between NE and DA are indicated. n = 3 independent cultures each. (D) Same as (B), except the sensors were expressed in cultured cortical neurons; n = 20 neurons from 3 cultures per group. (E) The on and off kinetics of the change in fluorescence were measured using high-speed line scan imaging of HEK293T cells expressing GRAB_NE2m_ or GRAB_NE2h_; τ_on_ was measured by fitting the rise in fluorescence upon rapid application of NE, and τ_off_ was measured by fitting the fluorescence decay upon application of the α2AR antagonist yohimbine (YO) in the continued presence of NE. E1 shows the experimental setup, including the line- scanning region and the pipette for rapid drug application. E2 and E3 show representative traces and the summary data, respectively; n ≥ 3 cells from 3 cultures per group. (F) Excitation (blue) and emission (green) spectra of GRAB_NE2m_ (left) and GRAB_NE2h_ (right) in the absence (dashed lines) and presence (solid lines) of 100 μM NE using one-photon imaging. (G) Summary of relative dose-dependent downstream G protein coupling of the wild-type α2 adrenergic receptor (WT-α2AR), GRAB_NE2m_, and GRAB_NE2h_ expressed in HEK293T cells measured using the luciferase complementation mini-G protein assay. n = 3 wells with ≥10^5^ cells each. (H) Summary of relative dose-dependent downstream β-arrestin coupling of WT-α2AR, GRAB_NE2m_, and GRAB_NE2h_ expressed in HEK293T cells measured using the Tango assay. n = 3 wells with ≥10^5^ cells each. The scale bars in (A) and (E) represent 20 μm; the scale bar in (D) represents 50 μm. Unless noted, summary data are presented as the mean ± SEM. \*\*\**p* < 0.001, \**p* < 0.05, and n.s., not significant (Student’s *t*-test and two-way ANOVA). See also Figure S1.

Next, we expressed the first-generation GRAB_NE1m_ sensor and our second-generation GRAB_NE2m_ and GRAB_NE2h_ sensors in HEK293T cells (Figure 1B) and found that applying 100 μM NE induced a peak ΔF/F_0_ of 230±9%, 381±23%, and 415±25%, respectively (Figure 1B2); in addition to their stronger response to NE, both GRAB_NE2m_ and GRAB_NE2h_ had higher maximum brightness compared to GRAB_NE1m_. In addition, both GRAB_NE2m_ and GRAB_NE2h_ retained the pharmacology of the parent α2AR receptor and do not respond to other neurochemicals, including the β2-adrenergic receptor agonist isoprenaline (ISO), acetylcholine (ACh), serotonin (5-HT), glutamate (Glu), γ-aminobutyric acid (GABA), adenosine (ADO), or histamine (HA); finally, the NE-induced response was blocked by the α2AR antagonist yohimbine (YO) but not the β2AR antagonist ICI-118,551 (ICI) (Figure S1). To test the performance of our GRAB_NE_ sensors in neurons, we expressed GRAB_NE2m_ and GRAB_NE2h_ in cultured cortical neurons. We generated dose-response curves for GRAB_NE2m_ and GRAB_NE2h_ and measured apparent affinity values of 320 nM and 78 nM, respectively (with 2-3 folds increase compared to GRAB_NE1m_), in response to NE, with significantly lower affinity for dopamine (350 μM and 110 μM, respectively) (Figure 1C). These resulted in an over 1000-fold selectivity to distinguish NE from DA for both GRAB_NE2m_ and GRAB_NE2h._ Both sensors’ peak ΔF/F_0_ were consistent with our results obtained using HEK293T cells (Figure 1D).

To determine whether the next-generation GRAB_NE_ sensors respond to NE with rapid kinetics, we locally puffed a saturating concentration of 10 μM NE onto HEK293T cells expressing either GRAB_NE2m_ or GRAB_NE2h_ and measured the change in fluorescence using high-speed line scan imaging (Figure 1E1). Fitting the rising phase of the fluorescence change using a single exponential function yielded average τ_on_ values of 0.12 s and 0.09 s for GRAB_NE2m_ and GRAB_NE2h_, respectively (Figure 1E2-3). We also fit the decrease in fluorescence following the addition of YO in the presence of NE and obtained average τ_off_ values of 1.72 s and 1.93 s for GRAB_NE2m_ and GRAB_NE2h_, respectively. The kinetics of GRAB_NE2m_ and GRAB_NE2h_ is a bit slower than GRAB_NE1m_^21^, possibly due to the higher affinity.

Next, we measured the spectral properties of GRAB_NE2m_ and GRAB_NE2h_ using one-photon excitation. We found that both sensors have excitation peaks at 410 nm and 500 nm, and an emission peak at 520 nm (Figure 1F), similar to the spectra of GFP and the calcium indicator GCaMP; thus, our sensors are compatible with various established imaging systems.

Because overexpressed GPCRs or their derivatives may induce downstream signaling, they have the potential to affect cellular physiology and may therefore be unsuitable for use in *in vivo* imaging. To rule out this possibility, we examined whether GRAB_NE2m_ and GRAB_NE2h_ induce downstream G protein and/or β-arrestin signaling using a luciferase complementation mini-G protein assay and the Tango assay, respectively (see Methods). We found that both sensors have negligible downstream coupling (Figure 1G and 1H), suggesting that overexpressing either GRAB_NE2m_ or GRAB_NE2h_ does not significantly affect cellular physiology.

### Detection of optogenetically evoked NE release in freely moving mice

Having shown that our next-generation GRAB_NE_ sensors have superior sensitivity, high specificity, rapid kinetics, and negligible downstream coupling *in vitro*, we then examined whether these sensors can report endogenous NE release *in vivo* when expressed in the locus coeruleus (LC) of TH-Cre mice together with the optogenetic actuator C1V1 linked to YFP (Figure 2A). For this experiment, we used spectrally resolved fiber photometry^23^ to simultaneously measure GRAB_NE_ and YFP. We found that optogenetic stimulation of LC-NE neurons elicited increases in GRAB_NE2m_ and GRAB_NE2h_ fluorescence in freely moving mice, but had no effect on YFP fluorescence (Figure 2B and 2C1). In addition, an intraperitoneal (i.p.) injection of the norepinephrine transporter (NET) inhibitor desipramine caused a progressive increase in the basal fluorescence of GRAB_NE2h_, reflecting an accumulation of extracellular NE and the high affinity of GRAB_NE2h_ for NE (Figure 2B); moreover, in the presence of desipramine the response induced by optogenetic stimulation was larger in magnitude and had slower decay kinetics (Figure 2C2). Conversely, an i.p. injection of the α2AR antagonist YO nearly abolished both the desipramine-induced increase in basal fluorescence and the optogenetic stimulation‒evoked increase in GRAB_NE2m_ and GRAB_NE2h_ fluorescence (Figure 2B and 2C3). In separate experiments, we injected the mice with either the selective dopamine transporter (DAT) inhibitor GBR-12909 followed by the D2R-specific antagonist eticlopride, which had no effect on basal fluorescence (data not shown) or the kinetics or magnitude of the optogenetically stimulated increase in GRAB_NE2m_ and GRAB_NE2h_ fluorescence (Figure 2C-E). Importantly, we found that GRAB_NE2m_ and GRAB_NE2h_ had an ∼17% and 24% increase in ΔF/F_0_, respectively, in response to a single train of light pulses (Figure 2D), a 2.4-3.9-fold improvement over the first-generation GRAB_NE1m_ sensor. These results suggest that our next-generation GRAB_NE_ sensors can reliably detect optogenetically evoked NE release in the LC of freely moving mice.

**Figure 2.**
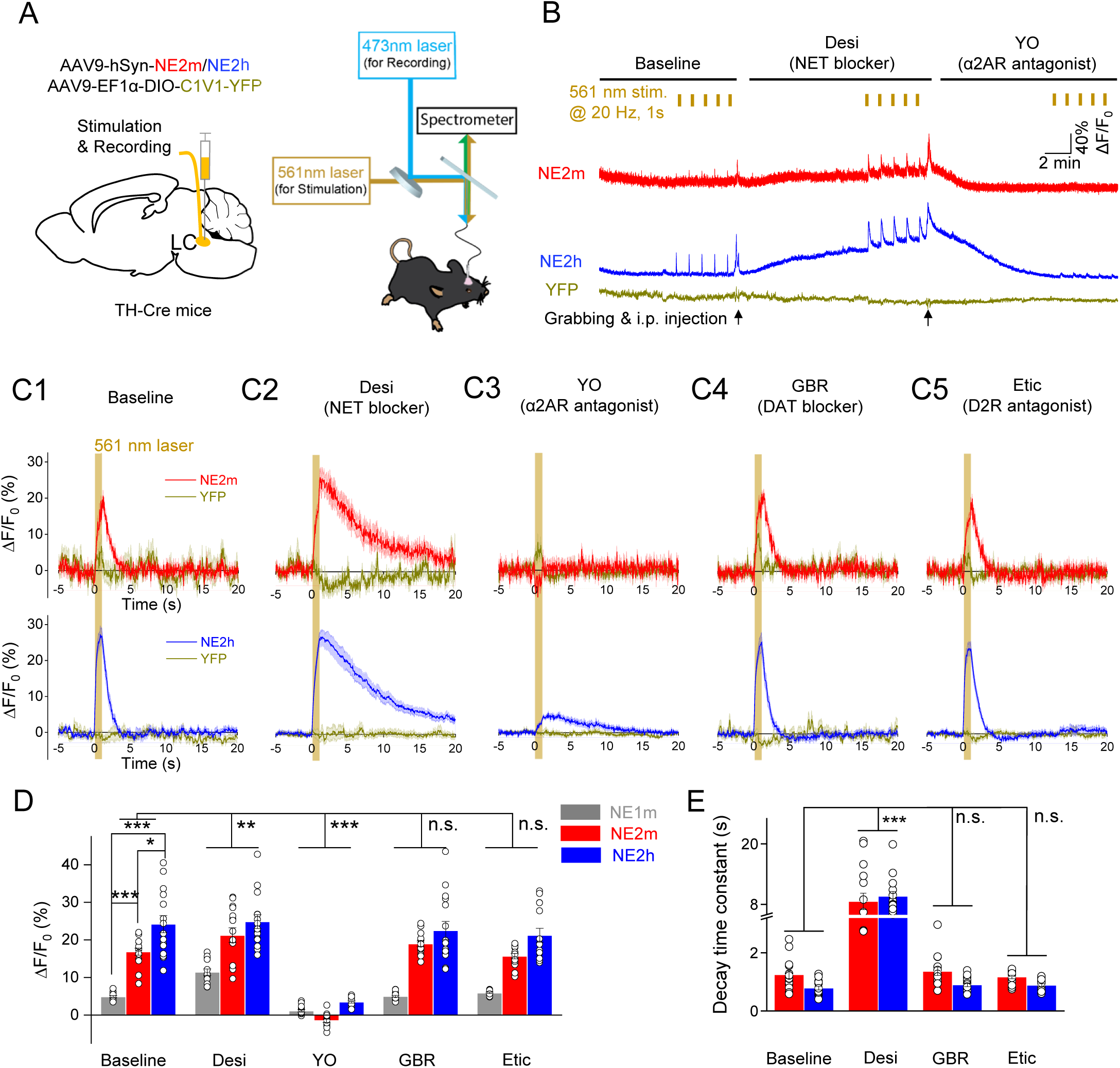
Detection of optogenetically evoked NE release in freely moving mice. (A) Experimental design depicting the strategy for expressing GRAB_NE2m_ and GRAB_NE2h_ and recording the change in fluorescence in response to optical stimulation of C1V1 in the locus coeruleus (LC). (B) Representative traces of optogenetically stimulated fluorescence change in GRAB_NE2m_ (red), GRAB_NE2h_ (blue), and YFP (olive, as a negative control) in the LC before (baseline, left), after an i.p. injection of the NE transporter (NET) blocker desipramine (Desi, 10 mg/kg, middle), and after an i.p. injection of the α2AR antagonist yohimbine (YO, 2 mg/kg, right). The vertical tick marks (yellow) indicate the optogenetic stimuli delivered at 20 Hz. (C-E) Average traces (C), summary of ΔF/F_0_ (D), and summary of decay time constants (E) of the change in fluorescence of GRAB_NE2m_ (top row in C, red) and GRAB_NE2h_ (bottom row in C, blue) in response to optical stimulation in the LC following treatment with the indicated compounds. Also shown in (C) are the fluorescence traces for YFP. The data for GRAB_NE1m_ in (D) and (E) were reproduced^21^ for comparison. n = 15 trials in 3 mice per group. GBR, GBR-12909; Etic, eticlopride. \*\*\**p* < 0.001, \*\**p* < 0.01, \**p* < 0.05, and n.s., not significant (two-way ANOVA).

### Next-generation NE sensors report behaviorally evoked NE release in vivo in response to stressful stimuli

The lateral hypothalamus (LH) is a target of the LC and has been shown to release NE during specific behaviors such as stress^21^. We therefore examined whether our next- generation GRAB_NE_ sensors can exhibit higher signals in measuring behaviorally evoked NE release in the LH of freely moving mice. We expressed GRAB_NE1m_, GRAB_NE2m_, or GRAB_NE2h_ in the LH of wild-type mice (Figure 3A and 3B) and then performed fiber photometry recordings during stress-inducing activities, including tail suspension (Figure 3C), forced swimming (Figure 3D), and hand presentation (Figure 3E). Consistent with previous reports^21^, all three stressors elicited an increase in GRAB_NE_ fluorescence. Moreover, both GRAB_NE2m_ and GRAB_NE2h_ had a larger response (up-to 3.7-fold) during tail suspension than GRAB_NE1m_ (Figure 3C). Interestingly, GRAB_NE2h_ had the largest response among all three sensors during both forced swimming and hand presentation (Figure 3C and 3E). In addition, an i.p. injection of the selective NET inhibitor atomoxetine induced a slow decay in the response to tail suspension, without significantly affecting peak ΔF/F_0_; in contrast, the α2AR antagonist YO significantly reduced the tail suspension‒evoked increase in GRAB_NE2m_ and GRAB_NE2h_ fluorescence (Figure 3F1 and 3G1). Finally, neither the selective DAT blocker GBR-12909 nor the D2R antagonist sulpiride affected the magnitude or kinetics of the response (Figure 3F2 and 3G2). These results indicate that our next-generation GRAB_NE_ sensors can be used to specifically monitor the release of endogenous NE in the LH in response to stress.

**Figure 3.**
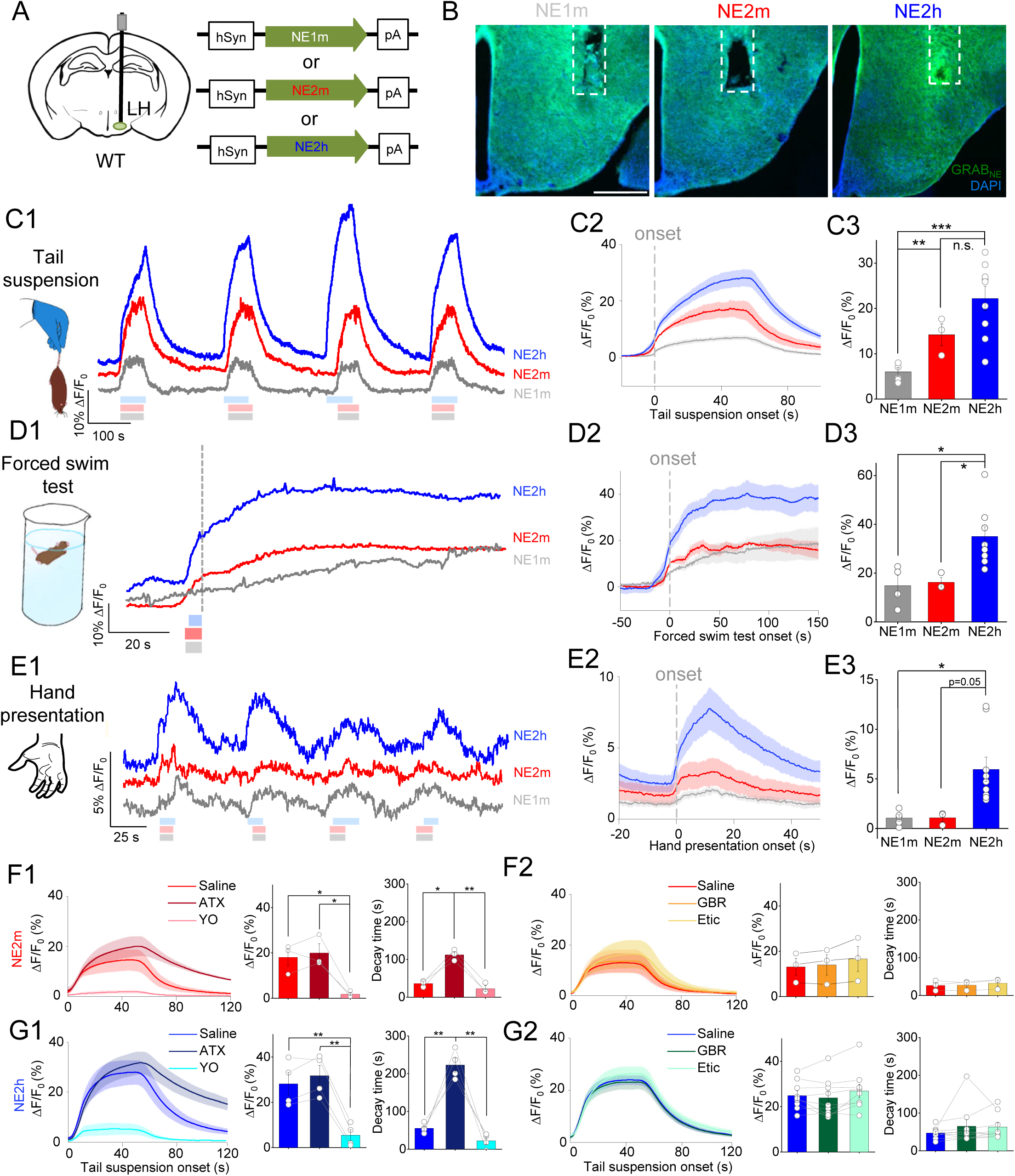
Next-generation NE Sensors report behaviorally evoked NE release *in vivo* in response to stressful stimuli. (A) Schematic diagram depicting the strategy for virus injection, fiber placement, and the recording site for GRAB_NE1m_, GRAB_NE2m_, or GRAB_NE2h_ in the lateral hypothalamus (LH). (B) Fluorescence images of brain sections of mice injected with virus expressing the indicated GRAB_NE_ sensors (green); the nuclei were counterstained with DAPI (blue). The position of the fiber is indicated by dashed white rectangles. Scale bar = 500 μm. (C-E) Representative traces (1), averaged per-stimulus histograms (2), and summary data (3) of GRAB_NE_ fluorescence (ΔF/F_0_) measured before, during, and after tail suspension (C), before and during forced swimming (D), and before, during, and after hand presentation (E); n ≥ 3 animals per group. The shaded bars in (C-E) indicate hand presentation to deliver respective stimuli. The grey dashed lines in (C-E) indicate the onset of respective stimuli. (F-G) Averaged per-stimulus histograms (left), summary data in GRAB_NE_ fluorescence (ΔF/F_0_) (middle), and post-test decay time (right) measured in mice expressing GRAB_NE2m_ (F) or GRAB_NE2h_ (G) in the LC during the tail suspension test 25 mins after an i.p. injection of saline (Sal), atomoxetine (ATX), yohimbine (YO), GBR-12909 (GBR), or eticlopride (Etic) as indicated; n ≥ 3 animals per group. \*\*\**p* < 0.001, \*\**p* < 0.01, *p < 0.05, and n.s., not significant (Student’s *t*-test).

### NE and calcium dynamics during the sleep-wake cycle

Genetically encoded GRAB_NE_ sensors can also be used to examine the spatiotemporal dynamics of NE release in the brain, which is a tightly regulated, complex process that can depend on a variety of factors such as the state of arousal and the activation of distinct brain regions. Moreover, previous studies suggested that specific brain regions may have either similar or distinct patterns of neurotransmitter release during the sleep-wake cycle^24-26^. To measure the dynamics of NE release in specific brain regions and determine whether this release is synchronized between brain regions, we utilized expressed GRAB_NE2m_ to simultaneously monitor NE levels in both the medial prefrontal cortex (mPFC) and the preoptic area of the hypothalamus (POA) (Figure 4A), two brain regions critically involved in regulating arousal and wakefulness. Meanwhile, we used electroencephalogram (EEG) and electromyogram (EMG) recordings to determine the animal’s sleep-wake state—i.e., awake, in NREM (non-rapid eye movement) sleep, or in REM (rapid eye movement) sleep. Dual-site continuous fiber photometry recording revealed that the changes in NE levels were closely synchronized between the mPFC and POA throughout the sleep-wake cycle (Figure 4B-D). Specifically, NE levels were relatively high during the wakefulness and NREM sleep, but low during REM sleep (Figure 4C), which is consistent with previous results^24, 25^. To analyze NE kinetics during the various state transitions, we calculated the t_50_ from each fluorescence trace and found similar kinetics between the mPFC and POA, with a rapid increase in NE release during the transition from REM sleep to the awake state (∼5 s) and from NREM sleep to the awake state (∼4 s), suggesting rapid NE release during arousal (Figure 4E-F). In contrast, the decrease in NE release was relatively slow during the transition from the awake state to NREM sleep (∼22 s) and from NREM sleep to REM sleep (∼30 s).

**Figure 4.**
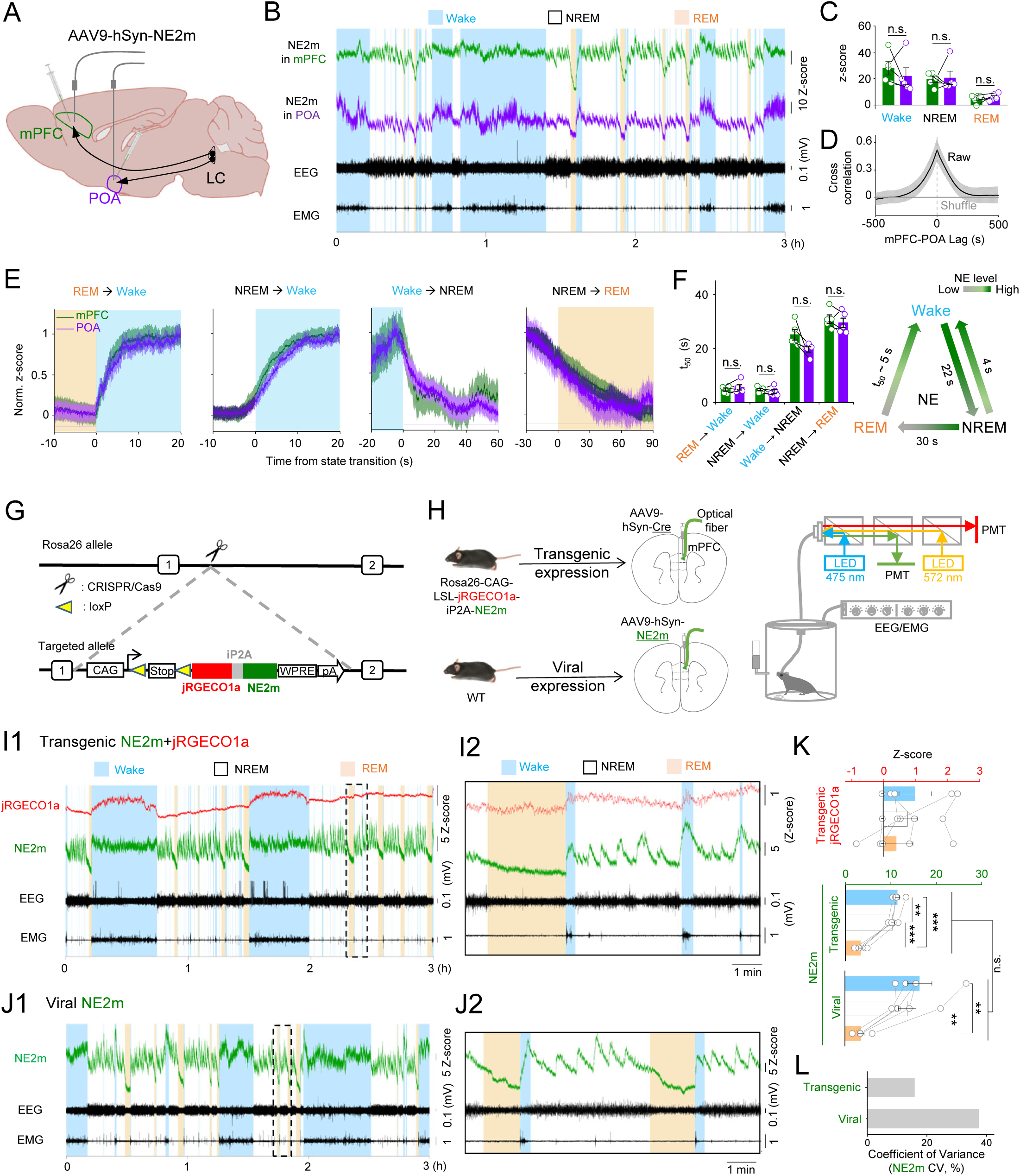
NE and calcium dynamics during the sleep-wake cycle. (A) Illustration depicting the strategy for virus injection and fiber placement for recording GRAB_NE2m_ fluorescence in both the medial prefrontal cortex (mPFC) and preoptic area of the hypothalamus (POA) during the sleep-wake cycle. The LC and its projections to the mPFC and POA are also indicated. (B-C) Representative traces of the GRAB_NE2m_ fluorescence signal (expressed as a *z*-score), EEG, and EMG recordings (B) and summary data of GRAB_NE2m_ fluorescence measured in mPFC and POA during the wake state, NREM sleep, and REM sleep (C). (D) Cross-correlation between GRAB_NE2m_ fluorescence measured in the mPFC and GRAB_NE2m_ fluorescence measured in the POA; also shown are the same raw data after being randomly shuffled. (E) Representative time courses of the GRAB_NE2m_ fluorescence signal measured in the mPFC and POA during the indicated transitions between the indicated sleep-wake states. (F) Summary data (left) and summary model (right) of the t_50_ values measured for each transition between the indicated sleep-wake states. (G) Strategy used to generate the dual-NECa transgenic knock-in mouse line expressing both GRAB_NE2m_ and jRGECO1a in the *Rosa26* locus. (H) Schematic illustration depicting the strategy used for virus injection and dual-color fiber photometry recording of GRAB_NE2m_ and jRGECO1a in the mPFC of dual-NECa transgenic mice (top) or wild-type (WT) mice (bottom) during the sleep-wake cycle. (I-K) Representative jRGECO1a, GRAB_NE2m_, EEG, and EMG traces (1), expanded traces (2) based on the dashed rectangle in (1), and summary (K) of the jRGECO1a and GRAB_NE2m_ signals measured in dual-NECa transgenic mice (I) or WT mice virally expressing GRAB_NE2m_ (J) during the awake state, NREM sleep, and REM sleep. (L) Coefficient of variation (CV) between the transgenic GRAB_NE2m_ and virally expressed GRAB_NE2m_ signals measured during the sleep-wake cycle. n = 5 animals per group. \*\*\**p* < 0.001, \*\**p* < 0.01, \**p* < 0.05, and n.s., not significant (two- way ANOVA for F, one-way ANOVA and Student’s *t*-test for K).

Although using virus injection to express genetically encoded sensors has several advantages, this approach also has several practical limitations, including the need for invasive surgery to inject the virus, limited region of delivery, variable levels of expression, and potential long-term cytotoxicity. To overcome these limitations, we generated a transgenic mouse line that expresses floxed GRAB_NE2m_ and jRGECO1a^27^—a red fluorescent calcium indicator—driven by the ubiquitous *CAG* promoter and targeted to the *Rosa26* locus^28–30^. Upon Cre expression, the cells in these mice express both GRAB_NE2m_ and jRGECO1a; these mice are referred to hereafter as dual-NECa mice (Figure 4G).

First, we virally expressed Cre in the mPFC of dual-NECa mice and used dual-color fiber photometry recording to measure both NE and calcium while monitoring the sleep-wake state using EEG and EMG (Figure 4H). We found that the GRAB_NE2m_ sensor expressed in the mPFC of our dual-NECa mice faithfully reported NE release throughout the sleep-wake cycle, consistent with previous reports^24, 25^. In addition, by measuring jRGECO1a fluorescence we observed relatively higher noradrenergic and calcium activities during the awake state, low noradrenergic and calcium activities during REM sleep, and distinct patterns of oscillatory NE release and relatively low calcium activity during NREM sleep (Figure 4I-J).

Importantly, neither the amplitude nor the kinetics of the NE signals measured in the dual-NECa transgenic mice differed significantly from the signals measured in mPFC neurons expressing GRAB_NE2m_ via AAV-mediated delivery (Figure 4I-K). Furthermore, the NE signals recorded in the dual-NECa mice had lower within-group variation than that of viral expression (Figure 4L). Taken together, these findings indicate that our dual-NECa transgenic mouse line is a useful tool to consistently report NE release and calcium dynamics simultaneously with spatial precision.

### Mesoscopic NE and calcium dynamics in dorsal cortex of awake mice

Another advantage of our dual-NECa mouse is that it can be crossed with established Cre-driver lines to express both GRAB_NE2m_ and jRGECO1a in specific cell types. Currently, approximately 500 Cre-driver lines are available from the Jackson Laboratory that express reporter genes either globally or in specific cell types and/or tissues throughout the central nervous system or periphery. We first crossed our dual-NECa reporter mouse with the CaMKIIα-Cre mouse (Figure 5A, top) in order to measure NE release and calcium dynamics specifically in excitatory neurons. The heterozygous mouse strain displays a healthy phenotype, with no significant abnormalities or defects in terms of growth, behavior, and reproduction. Noradrenergic neurons in the LC, which project to the entire brain and modulate a wide range of behaviors, including attention, stress, and cognition, have been reported to have high molecular and functional heterogeneity^31, 32^; thus, the pattern of NE release during these behaviors has remained poorly understood. Based on the one-photon spectra of GRAB_NE2m_ and jRGECO1a (Figure 5A, bottom), we performed cortex-wide two-channel imaging mesoscopy using 488-nm and 561-nm lasers to excite GRAB_NE2m_ and jRGECO1a, with an additional 405-nm laser signal used to correct for hemodynamic changes in the cortex^33^ (Figure 5B, see Methods).

**Figure 5.**
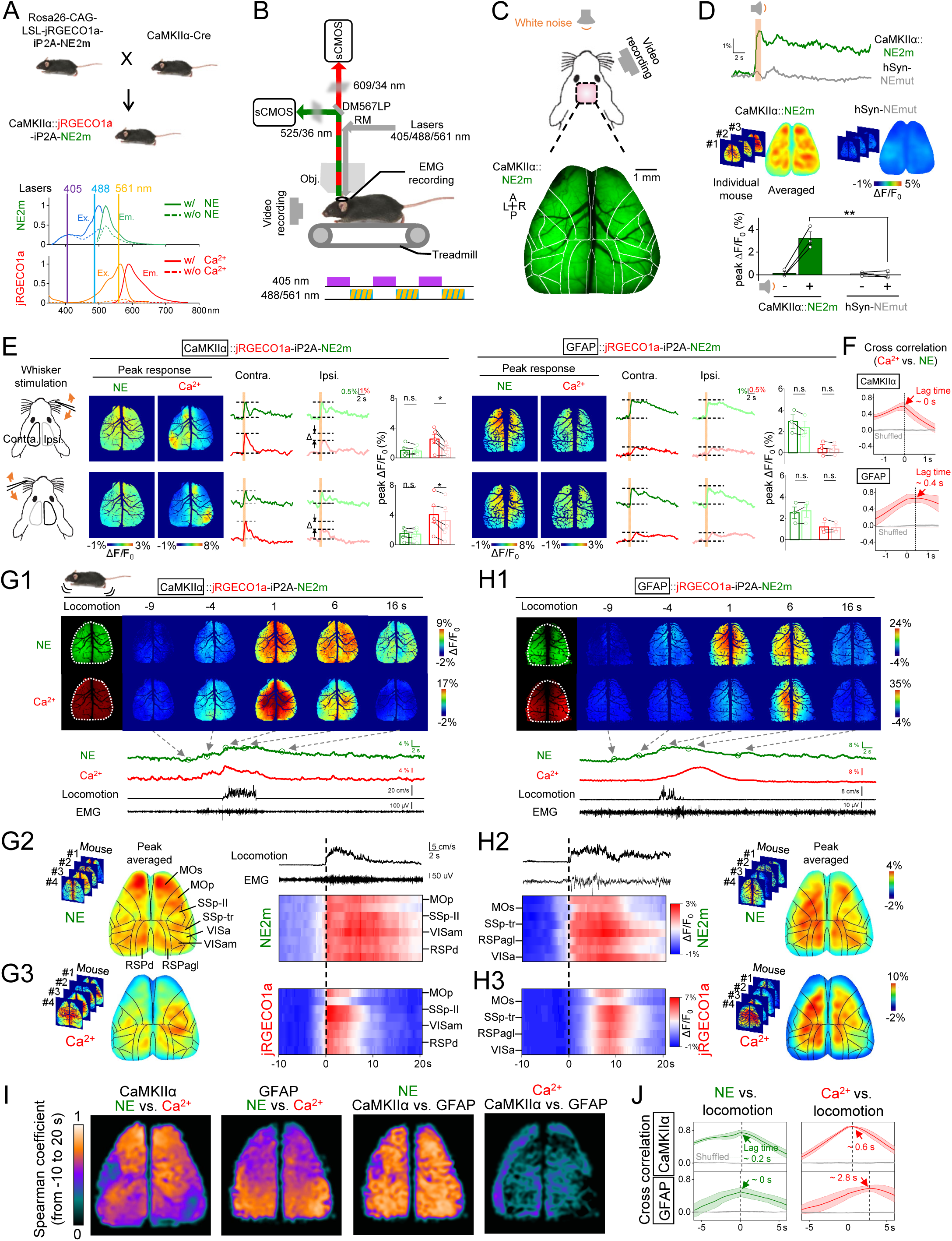
Mesoscopic NE and calcium dynamics in dorsal cortex of awake mice. (A) (Top) Schematic diagram depicting the strategy for generating CaMKIIα::NECa mice by crossing dual-NECa mice with CaMKIIα-Cre mice to drive the expression of GRAB_NE2m_ and jRGECO1a in excitatory neurons. (Bottom) One-photon excitation and emission spectra of GRAB_NE2m_ (in the absence and presence of ligand) and jRGECO1a^22^ (replotted from FPbase^35^); the three excitation lasers used for mesoscopic imaging are also indicated. (B) Schematic diagram depicting the dual-color mesoscopic imaging setup for recording GRAB_NE2m_ and jRGECO1a fluorescence in behaving mice. Excitation light alternated between green and red fluorescence imaging, and artifacts were corrected using 405-nm excitation. (C) Green and red fluorescence was measured using mesoscopic imaging through a 6 mm x 8 mm cranial window in CaMKIIα-Cre::NECa mice, with stimulation by a 1-s pulse of white noise. Shown below is an example image of GRAB_NE2m_ fluorescence. (D) Time course of the change in fluorescence intensity (top) and peak responses (bottom) measured in CaMKIIα::NECa mice and WT mice expressing the NE-insensitive GRAB_NEmut_ sensor (via virus injection at P0-P1; see Methods) before and immediately following audio stimulation. Peak response maps from individual mouse and averaged response map were shown. n = 3 animals per group. (E-F) Schematic illustration (left) of whisker stimulation delivered to CaMKIIα::NECa and GFAP::NECa mice co-expressing both jRGECO1a and GRAB_NE2m_ in excitatory neurons and astrocytes, respectively. Whisker stimuli were applied unilaterally to either the right (top row) or left (bottom row) side, and peak response images, representative traces, and the summary of relative peak ΔF/F_0_ measured in CaMKIIα::NECa (middle) and GFAP::NECa (right) mice are shown. The black and grey lines in the schematic illustration (left) indicate the ROIs used to analyze the representative traces and peak responses. Shown in (F) is the cross-correlation and time lag between the calcium and NE signals measured in response to bilateral whisker stimulation. The n = 5 animals per group. (G-H) Representative dual-color mesoscopic images (1, top) and traces (1, bottom) of GRAB_NE2m_ and jRGECO1a fluorescence measured in CaMKIIα::NECa (G) and GFAP::NECa (H) mice before, during, and after locomotion. Individual, averaged peak responses, and heatmaps of various cortical regions over time are shown for the GRAB_NE2m_ (2) and jRGECO1a (3) signals. The dashed white lines in (1) indicate the ROIs used to analyze the representative traces. n = 4 animals per group. (I) Cortex-wide Spearman coefficient measured between the NE and calcium signals in the CaMKIIα::NECa and GFAP::NECa mice (left two images) and between the CaMKIIα::NECa and GFAP::NECa mice for NE and calcium (right two images). (J) Cross-correlation and time lag between the NE and calcium signals and the onset of locomotion measured in CaMKIIα::NECa (top) and GFAP::NECa (bottom) mice. \*\**p* < 0.01, \**p* < 0.05 and n.s., not significant (Student’s *t*-test). See also Figures S2 and S3.

To confirm that the change in fluorescence measured using dual-color mesoscopy was specific for NE, we applied auditory stimuli to mice expressing either GRAB_NE2m_ or our previously reported NE-insensitive mutant sensor, GRAB_NEmut_^21^. We applied a 1-second pulse of white noise as the auditory stimulus and measured green fluorescence through a 6-mm x 8-mm D-shaped cranial window; we also used an infrared camera to record pupil size to confirm the mouse’s autonomic response to the auditory pulse (Figure 5C, top). We first verified that both GRAB_NE2m_ and GRAB_NEmut_ were expressed throughout the cerebral cortex (Figure 5C, bottom and Figure S2A). We then found that application of the auditory pulse induced a time-locked increase in fluorescence in the mice expressing GRAB_NE2m_, but had no effect in mice expressing GRAB_NEmut_ (Figure 5D). In contrast, the auditory pulse caused an increase in pupil diameter in both groups, indicating the presence of a general arousal response (Figure S2B). In addition, the relatively homogenous pattern of NE release in the cortex induced by the auditory stimulation (Figure 5D) is consistent with the reported distribution of LC fibers throughout the cortex^34^.

Next, to measure cell type‒specific noradrenergic and calcium signaling in response to tactile stimuli, we crossed our dual-NECa mouse line with mice expressing CaMKIIα-Cre or GFAP-Cre to drive the expression of both GRAB_NE2m_ and jRGECO1a in excitatory neurons and astrocytes, respectively; we then performed mesoscopic imaging and measured the change in NE and calcium in response to unilateral whisker stimulation (Figure 5E). In the CaMKIIα::NECa mice, we observed a time-locked global increase in GRAB_NE2m_ fluorescence throughout the dorsal cortex, while the calcium signal increased only in the contralateral hemisphere, consistent with thalamocortical projections (Figure 5E). In addition, bilateral whisker stimulation induced a symmetrical concurrent increase in both NE and calcium (Figure 5F and Figure S3A1). In contrast, we observed a similar global increase in GRAB_NE2m_ fluorescence in the GFAP::NECa mice, but a relatively small and delayed calcium signal increase during unilateral and bilateral whisker stimulation (Figure 5E, 5F, and Figure S3A2).

As a further test of the effect of sensory stimuli on NE and calcium signaling in different cell types, we delivered either binocular or monocular visual stimuli to these mice. In the CaMKIIα::NECa mice, monocular visual stimulation induced an increase in calcium in the contralateral visual cortex (Figure S3C). Interestingly, however, binocular visual stimulation induced a small but measurable global increase in GRAB_NE2m_ fluorescence throughout the cortex, while monocular stimulation had no effect on either hemisphere. In contrast, visual stimulation had no effect on either NE or calcium signaling in the GFAP::NECa mice (Figure S3B). Thus, dual-color mesoscopic imaging of our dual-NECa transgenic mice revealed cell type‒specific differences in the spatiotemporal patterns of NE and calcium signaling in response to distinct sensory inputs, providing valuable insights into the underlying neural circuitry.

Finally, we examined NE and calcium signaling in response to spontaneous locomotor activity using the EMG data and the speed of the linear treadmill as a measure of the onset and duration of locomotion, respectively. In the CaMKIIα::NECa mice, we found that both the NE and calcium signals increased in the dorsal cortex with increased locomotor activity (Figure 5G1); similar results were obtained in the GFAP::NECa mice (Figure 5H1). We then aligned and averaged the peak response images of both the NE and calcium signals obtained from each mouse during locomotion, and then segmented the dorsal cortex into distinct brain regions using the Allen Brain Atlas (Figure 5G2-H3). We found that in both the CaMKIIα::NECa and GFAP::NECa mice NE was released globally, with a high Spearman coefficient throughout the cortex (Figure 5I). In contrast, the calcium signal increased in the sensorimotor cortex in “hotspots” in the CaMKIIα::NECa mice, while the calcium signal increased globally in the GFAP::NECa mice with an ∼2.75 second delay following the onset of locomotion (Figure 5J). These results shed new light on the intricate interplay between NE and calcium signaling in the brain during distinct behaviors.

## Discussion

Here, we developed an optimized set of next-generation GRAB_NE_ sensors with an increased response, sensitivity, and molecular selectivity for NE. We then used these new sensors to detect optogenetically and behaviorally triggered NE release in the locus coeruleus and lateral hypothalamus of freely moving mice. In addition, we developed a novel transgenic mouse line expressing both GRAB_NE2m_ and the calcium sensor jRGECO1a in specific cell types and performed simultaneous dual-color recording and cell type‒specific spatiotemporal imaging of NE and calcium signaling during the sleep-wake cycle, sensory processing, and locomotion in behaving mice.

Given the structural similarity and widespread patterns of NE and DA throughout the brain, distinguishing these two monoamines is essential when performing *in vivo* behavioral studies. Moreover, because human noradrenergic receptors respond to both NE and DA, our goal is to increase the sensitivity of our GRAB_NE_ sensors to NE while reducing their response to DA. Using cell-based screening, we identified specific combinations of mutations that increased the sensors’ sensitivity to NE without compromising their selectivity, underscoring the power of high-throughput screening in navigating complex chemical spaces. Moreover, consistent with our *in vitro* results, our next-generation GRAB_NE_ sensors produce robust *in vivo* signals in response to both optogenetic stimulation and behavioral events.

Importantly, our dual-NECa transgenic mouse allows for the simultaneous monitoring of NE and calcium signaling in specific cell types and brain regions with high spatiotemporal resolution during a wide variety of physiological conditions and stimuli. Although the sensors’ expression levels are presumably lower compared to AAV-mediated expression, the dual-NECa mice revealed similar changes in NE dynamics throughout the sleep-wake cycle, encompassing the signal amplitude and kinetics during state transitions. Moreover, a clear advantage of the dual-NECa mice is that the signals obtained have considerably lower within-group variance compared to viral expression, reflecting the reliability and high replicability of using the dual-NECa mouse to image NE and calcium signaling. Furthermore, this transgenic mouse line can be used to express the NE and calcium sensors in virtually any cell type and/or tissue by crossing with specific Cre-reporter mice.

Finally, using dual-color mesoscopic imaging of dual-NECa mice, we observed global versus “hotspot” patterns of NE release and cell type‒specific calcium signaling during distinct sensory processing events and locomotion. Thus, integrating cortex-wide imaging with our dual-NECa reporter mice offers a unique opportunity to examine NE and calcium signaling on a large scale with both cell type and molecular specificity in a wide range of physiological and pathophysiological contexts.

## Acknowledgments

This work was supported by the National Key R&D Program of China (2021YFF0502904 to J.F.), the National Natural Science Foundation of China (31925017 and 31871087 to Y.L.). The study was also supported by grants from the NIH BRAIN Initiative (U01NS120824 to Y.L.), the Feng Foundation of Biomedical Research, the Clement and Xinxin Foundation, the Peking-Tsinghua Center for Life Sciences, the State Key Laboratory of Membrane Biology at Peking University School of Life Sciences, and the New Cornerstone Science Foundation through the New Cornerstone Investigator Program and the XPLORER PRIZE (to Y.L.), the Leon Levy Neuroscience Fellowship and NIMH K99MH127295 grants (J.E.L), U01NS113358 and U01NS103558 (D.L.)., and the Intramural Research Program of the NIH/NIEHS of the United States (1ZIAES103310 to G.C.). We thank Xiaoguang Lei at PKU-CLS and the National Center for Protein Sciences at Peking University in Beijing, China, for their support and assistance with the Opera Phenix high-content screening system and imaging platform. We thank Yueyue Yu for help with maintaining the transgenic mice.

## Author contributions

Y.L. supervised the project. J.F. and Y.L. designed the study. J.F. and H.W. performed the experiments related to sensor optimization and characterization in cultured HEK293T cells and neurons. J.F. designed and constructed the dual-NECa transgenic mouse line. H.D. and X.M. performed the experiments related to the sleep-wake cycle. J.Z., and G.C. designed and performed the optogenetic stimulation experiments. J.L. and D.L. designed and performed the experiments involving behavior-related recording. J.F. and F.D. designed and performed the mesoscopic imaging experiments with help from H.X., C.Z. All authors contributed to the data interpretation and data analysis. J.F. and Y. L. wrote the manuscript with input from all other authors.

## STAR Methods

### EXPERIMENTAL MODEL AND SUBJECT DETAILS

#### Cell lines

HEK293T cells (cat. no. CRL-3216) were obtained from ATCC, cultured, and verified by their morphology and growth curve. The HTLA cells used in the Tango assay stably express a tTA-dependent luciferase reporter and a β-arrestin2-TEV fusion gene, and were generously provided by Bryan L. Roth^36^. All cell lines were cultured at 37°C in DMEM (Gibco) supplemented with 10% fetal bovine serum (Gibco) and 1% penicillin-streptomycin (Gibco) in humidified air containing 5% CO_2_.

#### Primary cell cultures

Postnatal day 0 (P0) Sprague-Dawley rat pups of both sexes, randomly selected from Beijing Vital River, were used to isolate cortical neurons. In brief, the brains were removed, the cortex dissected, neurons were dissociated in 0.25% Trypsin-EDTA (Gibco) was used to dissociate the neurons. The cells were subsequently plated on 12-mm glass coverslips coated with poly-D-lysine (Sigma-Aldrich) and cultured at 37°C in neurobasal medium (Gibco) supplemented with 2% B-27, 1% GlutaMax, and 1% penicillin-streptomycin (Gibco) in humidified air containing 5% CO_2_.

#### Mice/rats

All animal experiments were performed in accordance with the US National Institutes of Health guidelines for the care and use of laboratory animals, and were approved by the respective Animal Care and Use Committees at Peking University, New York University, and the US National Institute of Environmental Health Sciences. All animals were housed in pairs or as families in a temperature-controlled room with a 12-hour light-dark cycle (lights on from 10 am to 10 pm) with *ad libitum* access to food and water. The *in vivo* experiments were performed on adult (2-12 months of age) mice of both sexes.

TH-Cre mice (MMRRC_031029-UCD) were obtained from MMRRC. Dual-NECa transgenic mice were generated with help of Biocytogen Pharmaceuticals Co., Ltd. (Beijing, China) as follows. We designed and developed a floxed transgenic mouse line (dual-NECa, EGE-XWY-076) expressing GRAB_NE2m_-iP2A-jRGECO1a by targeting the *Rosa26* locus^30^. We first constructed a targeting vector containing the *CAG* promoter followed by the GRAB_NE2m_ and jRGECO1a coding sequences, separated by an improved P2A self- cleaving peptide^37^ to allow for independent expression of the two proteins. We then used CRISPR/Cas9-mediated homology-directed repair (HDR) to insert the targeting vector into the *Rosa26* locus of mouse embryonic stem cells. Successful targeting was confirmed via PCR-based screening and sequencing of the targeted genomic region. Next, the genetically modified embryonic stem cells were injected into eight-cell stage embryos to generate chimeric mice. The chimeric mice were then mated with wild-type mice to obtain germline transmission of the targeted allele. The resulting dual-NECa transgenic mouse line stably expressed both the green fluorescent GRAB_NE2m_ sensor and the red calcium indicator jRGECO1a under the control of the *CAG* promoter at the *Rosa26* locus upon excision of the floxed stop codon by Cre recombinase. CaMKIIα-Cre (005359; JAX) and GFAP-Cre (024098; JAX) were used in this study to further drive the expression of GRAB_NE2m_ and jRGECO1a.

### METHOD DETAILS

#### Molecular cloning

In this study, the molecular clones were generated using Gibson assembly. The DNA fragments were amplified with primers containing 25--30-bp overlap, and the cloning enzymes included T5-exonuclease, Phusion DNA polymerase, and Taq ligase. Sanger sequencing was used to confirm the sequence of all clones. The pDisplay vector with an upstream IgK leader sequence upstream and a downstream IRES-mCherry-CAAX cassette was used to clone all cDNAs encoding the GRAB_NE_ sensors, providing cell membrane targeting and labeling. For sensor optimization, amino acids were randomly mutated using PCR amplification with NNB codons at the target sites. The pAAV vector containing the human *Synapsin* promoter was used to clone express the GRAB_NE_ sensors or GRAB_NEmut_ in neurons. For luciferase complementation assay, the GRAB_NE_-SmBit and a2AR-SmBit constructs were modified from β2AR-SmBit, and the LgBit-mGsi was a gift from Nevin A. Lambert.

#### Expression of GRAB_NE_ sensors in cultured cells and *in vivo*

GRAB_NE_ sensors were expressed in HEK293T cells and cultured rat cortical neurons as previously reported^21^.

For *in vivo* virus-mediated expression, adult mice were anesthetized with either an i.p. injection of 2,2,2-tribromoethanol (Avertin, 500 mg/kg body weight, Sigma-Aldrich) or 1.5% isoflurane by inhalation, 2% lidocaine hydrochloride was injected subcutaneously under the scalp. the mice were then placed in a stereotaxic frame (RWD Life Science). Small craniotomy holes were prepared in the skull for virus injection.

In Figure 2, AAVs expressing hSyn-GRAB_NE2m/NE2h_ and Ef1a-DIO-C1V1-YFP^38^ (Vigene, 1×10^13^ titer genomic copies per ml) were injected into the LC (AP: -5.45 mm relative to Bregma; ML: ± 1.25 mm relative to Bregma; DV: 2.25 mm below the dura) of TH-Cre mice at a rate of 100 nl/min and in a volume of 500 nl. Four weeks after virus injection, we implanted multi-mode optical fiber probes (105/125 μm core/cladding) into the LC (AP: -5.45 mm relative to Bregma; ML: ± 0.85 mm relative to Bregma; DV: 3.5 mm below the dura).

In Figure 3, AAVs expressing GRAB_NE1m_, GRAB_NE2m_, and GRAB_NE2h_ (Vigene, 1×10^13^ titer genomic copies per ml) were unilaterally injected into the lateral hypothalamus (AP: -1.7 mm relative to Bregma; ML: +0.90 mm relative to Bregma; DV: 6.05 mm below the dura) of wild-type C57BL/6 mice at a rate of 10 nl/min and in a volume of 100 nl. A 400-μm optic fiber (Thorlabs, BFH48-400) housed in a ceramic ferrule (Thorlabs, SFLC440-10) was implanted 0.2 mm above the injection site. The experiments were performed three weeks after virus injection.

For the experiments in Figure 4, a fine glass pipette and a micro-syringe pump (Nanoliter 2010 injector, World Precision Instruments) were used to microinject approximately 300 nl of AAV9-hSyn-NE2m or AAV9-hSyn-Cre virus (Vigene, 1×10^13^ titer genomic copies per ml) into the mPFC (AP: +1.9 mm relative to Bregma, ML: -0.3 mm relative to Bregma, DV: 1.9 mm below the dura) and/or POA (AP: 0 mm relative to Bregma, ML: -0.6 mm relative to Bregma, DV: 4.9mm below the dura) at a rate of 30 nl/min.

For experiments in Figure 5, we crossed homozygous of the floxed dual-NECa transgenic mice with CaMKIIα-Cre (005359; JAX) or GFAP-Cre (024098; JAX) to obtain CaMKIIα::NECa and GFAP::NECa offspring, respectively. To achieve widespread expression of GRAB_NEmut_ through the entire cortex, we utilized a method previously described^39^ in which 4 μl of AAV9-hSyn-NEmut virus (Vigene, 1×10^13^ titer genomic copies per ml) was bilaterally injected into the transverse sinus of P0-P1 C57BL/6 mouse pups at a rate of 1.2 μl/min.

#### Fluorescence imaging of HEK293T cells and cultured neurons

To visualize cells expressing GRAB_NE_ sensors, we used either an inverted Ti-E A1 confocal microscope (Nikon) equipped with a 10x/0.45 NA (numerical aperture) objective, a 20x/0.75 NA objective, a 40x/1.35 NA oil-immersion objective, a 488-nm laser, and a 561- nm laser or an Opera Phenix high-content screening system (PerkinElmer) equipped with a 20x/0.4 NA objective, a 40x/1.1 NA water-immersion objective, a 488-nm laser, and a 561-nm laser. For confocal microscopy, the GFP signal was collected using a 525/50-nm emission filter combined with the 488-nm laser, while the RFP signal was collected using a 595/50-nm emission filter combined with the 561-nm laser. For the Opera Phenix system, the GFP and RFP signals were collected using a 525/50-nm and 600/30-nm emission filter, respectively. To calibrate the fluorescence signal produced by the green fluorescent GRAB_NE_ sensors, we used the GFP/RFP ratio. The dose-dependent response and on and off kinetics were determined as previously described^21^.

#### Measurements of spectra

HEK293T cells expressing GRAB_NE2m_ or GRAB_NE2h_ were harvested and transferred to a 384-well plate. Excitation and emission spectra were measured at 5-nm increments with a 20-nm bandwidth using a Safire2 multi-mode plate reader (TECAN) in the presence or absence of 10 μM NE. Control cells not expressing a sensor were used to obtain background fluorescence for subtraction.

#### Tango assay

HTLA cells expressing the wild-type α2AR, GRAB_NE2m_, or GRAB_NE2h_ were exposed to varying concentrations of NE (ranging from 0.1 nM to 10 μM) and cultured for 12 hours to allow luciferase gene expression. Luminescence was then measured using a VICTOR X5 multilabel plate reader (PerkinElmer) after adding Furimazine (NanoLuc Luciferase Assay, Promega) to a final concentration of 5 mM.

#### Luciferase complementation assay

The luciferase complementation assay was performed as described previously^40^. Forty- eight hours after transfection, the cells were washed with phosphate-buffered saline and transferred to opaque 96-well plates containing diluted NE solutions ranging from 1 nM to 100 μM. Luminescence was measured using Nluc after adding Furimazine (NanoLuc Luciferase Assay, Promega) to each well.

#### Fiber photometry recordings in freely moving mice during optical stimulation

In Figure 2, fiber photometry recording in the LC was performed using a 473-nm laser, which produced an output power of 25 μW at the end of the fiber. The resulting emission spectra were analyzed using a linear unmixing algorithm (https://www.niehs.nih.gov/research/atniehs/labs/ln/pi/iv/tools/index.cfm). The coefficients from the unmixing algorithm represent the fluorescence intensities of various fluorophores^23^. To evoke C1V1-mediated NE release, pulse trains (10-ms pulses at 20 Hz for 1 s) were delivered to the LC using a 561-nm laser with an output power of 9.9 mW at the end of the fiber.

#### Fiber photometry recordings in mice during behavioral testing

For the fiber photometry recordings in Figure 3, GRAB_NE_ sensors were excited using a 400-Hz sinusoidal blue LED light (30 mW; M470F1 driven by an LEDD1B driver; both from Thorlabs), which was bandpass filtered (passing band: 472 ± 15 nm, Semrock, FF02- 472/30-25) and transmitted to the brain. The emission light traveled back through the same optic fiber, through a bandpass filter (passing band: 534 ± 25 nm, Semrock, FF01-535/50), and was recorded using a Femtowatt Silicon Photoreceiver connected to an RZ5 real-time processor (Tucker-Davis Technologies). A custom-written program was used to extract the 400-Hz signals in real-time and determine the intensity of the GRAB_NE_ fluorescence signal.

All behavioral tests were performed at least 1 hour after the onset of the dark cycle. For the tail suspension test, each mouse was lifted gently off the bottom of its cage six times for 60 seconds each, with a minimum of 1 min between each lift. In the forced swimming test, the mouse was gently placed in a 1000-ml conical flask filled with lukewarm water and then removed after 4-6 min. the mouse was then gently dried with paper towels and placed on a heating pad inside its home cage. No aggressive behavior was observed during the test. All videos were recorded at 25 frames per second and manually annotated frame-by-frame using a custom MATLAB program (MathWorks)^41^.

#### Fiber photometry recordings and polysomnographic recordings during the sleep-wake cycle

To measure the fluorescence signals in Figure 4, a 200-μm optical fiber cannula (Fiber core: 200 μm; numerical aperture: 0.37; Inper, Zhejiang, China) was implanted 0.1 mm above the virus injection site and fixed to the skull using dental cement.

To monitor the animal’s sleep-wake state, EEG electrodes were implanted into the craniotomy holes above the frontal cortex and visual cortex, and EMG wires were placed in the trapezius muscles on both sides. The electrodes were connected to a microconnector and fixed to the skull using dental cement. The microconnector was connected via a flexible cable and attached to an electric slip ring, allowing the mouse to move freely. The cortical EEG and neck EMG signals were amplified (NL104A, Digitimer), filtered (NL125/6, Digitimer), digitized using a Power1401 digitizer (Cambridge Electronic Design Ltd.), and recorded using Spike2 software (Cambridge Electronic Design Ltd.) at a sampling rate of 1000 Hz.

A fiber photometry system (Thinker Tech, Nanjing, China) was used to record the fluorescence signals in freely moving mice. Blue (473-nm) and yellow (580-nm) LED lights (Cree LED) were bandpass filtered (470/25 nm, model 65-144 and 572/28 nm, model 84100, Edmund Optics), reflected by a 495-nm long-pass dichroic mirror (model 67-069, Edmund Optics) and a multi-band filter (model 87-282, Edmund Optics) dichroic mirror, and then focused using a 20x objective lens (Olympus). An optical fiber guided the light between the commutator and the implanted optical fiber cannula. The excitation light power at the tip of the optical fiber was adjusted to 20-30 μW in order to minimize photobleaching and was delivered at 100 Hz with a 5-ms pulse duration. Green fluorescence was bandpass filtered (525/39 nm, model MF525-39, Thorlabs), red fluorescence was bandpass filtered (615/20 nm, model 87753, Edmund Optics), and the resulting emissions were collected using a photomultiplier tube (model H10721-210, Hamamatsu). The current output from the photomultiplier tube was converted to a voltage signal using an amplifier (model C7319, Hamamatsu) and passed through a low-pass filter. The analog voltage signals were then digitized using an acquisition card (National Instruments). Photometry signals and polysomnographic recordings were aligned based on a TTL signal. To minimize autofluorescence of the optical fiber, the recording fiber was photobleached using a high-power LED before recording. Background autofluorescence was subtracted from the recorded signals during subsequent analysis.

#### Mesoscopic *in vivo* imaging

The surgery to prepare the imaging window and implant the EMG electrodes was performed on CaMKIIα::NECa, GFAP::NECa, or wild-type mice expressing GRAB_NEmut_. Anesthesia was induced with an i.p. injection of 2,2,2-tribromoethanol (Avertin, 500 mg per kg) and maintained with 1% isoflurane. The mouse was then fixed in a stereotaxic frame, and 2% lidocaine hydrochloride was injected under the scalp. To protect the corneas, erythromycin ophthalmic ointment was applied to both eyes. The scalp and underlying muscles were carefully removed to expose the skull, and the majority of the skull above the dorsal cortex was replaced with a custom-made coverslip to create an optical window. EMG electrodes were implanted as described above, and the mice were given at least 7 days to recover, followed by an additional 3 days to habituate to the head fixation before imaging.

Mesoscopic imaging was performed using a customized dual-color macroscope equipped with a 2x/0.5 NA objective lens (Olympus, MVPLAPO2XC), two 1x/0.25 NA tube lenses (Olympus, MVPLAPO1X), and two sCMOS cameras (Andor, Zyla 4.2 Plus, 2,048×2,048 pixels, 16-bit). A multi-line fiber-coupled laser system (Changchun New Industries Optoelectronics Tech. Co., Ltd., RGB-405/488/561/642nm-220mW-CC32594) generated three excitation wavelengths (405 nm, 488 nm, and 561 nm). Emission light was passed through a long-pass dichroic mirror (Thorlabs, DMLP567L) and either a 525/36-nm or 609/34-nm emission filter (Chroma) and captured by the sCMOS cameras. Both the excitation laser and the camera exposure were triggered by an Arduino board (Uno) using custom-written programs. Dual-color imaging was performed using alternating illumination between the 405-nm laser and the 488-nm or/and 561-nm laser. Images were acquired using Micro-Manager 2.0 at 512×512-pixel resolution at a rate of 5 Hz with 40-ms exposure.

During imaging, the mice were head-fixed but could run freely on a linear treadmill. A near-infrared camera with an infrared LED was used to record the mouse’s behavior and pupil size. For auditory stimulation, 1 sec of 70-dB white noise was generated using a RZ6 Multi I/O Processor (Tucker-Davis Technologies) and delivered via a magnetic speaker. For whisker stimulation, a 1-sec pendular stick was delivered to the mouse whisker either unilaterally or bilaterally. For visual stimulation, 50-ms of a flashing LED light was delivered to the mouse eye either unilaterally or bilaterally. Locomotion activity was recorded using the encoder in the treadmill.

#### Quantification and statistical analysis

For the imaging experiments using cultured HEK293T cells and primary neurons, fluorescence intensity was first quantified using ImageJ software (National Institutes of Health) or Harmony software (PerkinElmer, Inc.) for and then analyzed using a custom-written MATLAB script (MathWorks) or Origin Pro (OriginLab).

The photometry data were analyzed using a custom program written in MATLAB. To calculate ΔF/F_0_, baseline values were measured during REM sleep with no apparent fluctuations. To compare the change in fluorescence between animals, the *z*-score‒ transformed ΔF/F_0_ was normalized using the standard deviation of the baseline signals.

EEG and EMG recordings were used to determine the animal’s sleep-wake state. In brief, the EEG and EMG data were filtered at 0.5-100 Hz and 30-500 Hz, respectively, and semi-automatically scored off-line in 4-s epochs of wakefulness, REM sleep, and NREM sleep using AccuSleep (https://github.com/zekebarger/AccuSleep)^42^; the defined sleep-wake states were confirmed by visual examination and corrected if necessary. Wakefulness was defined as desynchronized low-amplitude EEG activity and high-amplitude EMG activity with phasic bursts. NREM sleep was defined as synchronized EEG activity with high-amplitude delta rhythm (0.5-4 Hz) and low EMG activity. REM sleep was defined as a pronounced theta rhythm (6-10 Hz) and low EMG activity. EEG spectral analysis was estimated using a short-time fast Fourier transform (FFT).

For the mesoscopic imaging data, raw images acquired from each camera were calibrated to ensure uniformity across the imaging region, and movement-related artifacts were corrected using the motion-correction algorithm NoRMCorre^43^. The corrected image stack with a size of 512 × 512 pixels was downsampled by a factor of 0.5 to 256 × 256 pixels for further analysis. For dual-color imaging, the red-channel images were registered to the green-channel images by performing an automated transformation using the “similarity” mode of the MATLAB function “imregtform”. The same transformation was then applied to all red-channel images to align them with their corresponding green-channel images. The resulting image stack was saved as a binary file to facilitate the input and output of large files. A mask was created to exclude background and blood vessel pixels from the corrected image stack using the machine learning‒based ImageJ plugin Trainable Weka Segmentation (v3.3.2); these minimized artifacts caused by blood vessel constriction and dilation. To correct the effects of hemodynamics on fluorescence^44, 45^, we performed a pixel-by-pixel correction based on a linear regression of the ligand-dependent signals (excited by 488-nm or 561-nm light) against the ligand-independent signals (excited by 405-nm light) for both GRAB_NE2m_ and jRGECO1a based on their respective spectra.

Baseline images were smoothed using a Gaussian filter (σ=2), and linear regression was performed for each pixel by regressing the baseline fluorescence intensity of the 405-nm‒ excited channel onto the 488-nm or 561-nm signal. The regression coefficient was then used to rescale the 405-nm channel, which was then subtracted from the 488-nm or 561- nm signal. The corrected signal was added to the averaged rescaled 405-nm channel signal to avoid negative values. The response of each pixel was calculated using the following equation: ΔF/F_0_ = (F−F_0_)/F_0_, where F_0_ is defined as the average baseline fluorescence intensity.

We registered the mean fluorescence image to a 2D projection of the Allen Common Coordinate Framework v3 (CCFv3) using four manually identified anatomical landmarks, including the left, center, and right points in the boundary between the anterior cortex and the olfactory bulbs, and the medial point at the base of the retrosplenial cortex. To analyze the time course of the response in a specific brain region, we calculated the average ΔF/F_0_ value for all available pixels within that region. To align and average the responses across the entire cortex from multiple mice, we developed a custom script to first register the peak response image for each individual mouse to the Allen CCFv3 and then averaged the images, preserving only the intersection pixels.

### DATA AND SOFTWARE AVAILABILITY

The custom-written MATLAB programs used in this study will be provided upon request to the corresponding author.

## Supplemental figure legends

**Figure S1.**
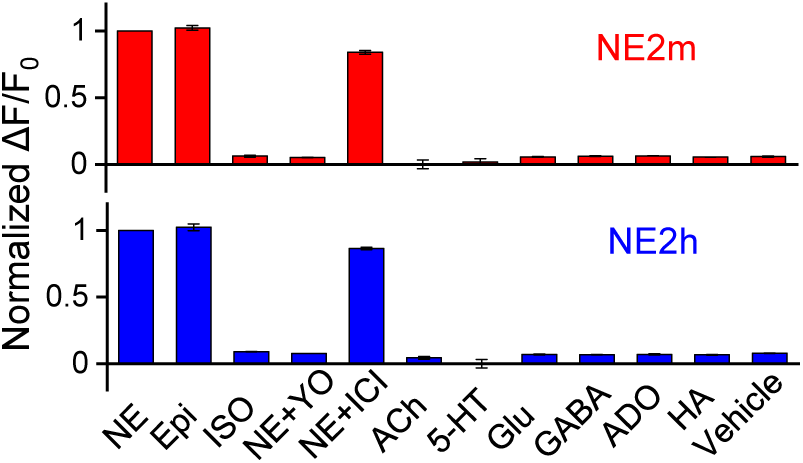
Selectivity of next-generation GRAB_NE_ sensors (related to Figure 1). Normalized changes in the fluorescence intensity of GRAB_NE2m_ (top) and GRAB_NE2h_ (bottom) in response to application of the indicated molecules (applied at 10 μM), expressed relative to NE. NE, norepinephrine; Epi, epinephrine; ISO, isoprenaline; YO, yohimbine; ICI, ICI-118,551; ACh, acetylcholine; 5-HT, 5-hydroxytryptamine (serotonin); Glu, glutamate; GABA, γ-aminobutyric acid; ADO, adenosine; HA, histamine.

**Figure S2.**
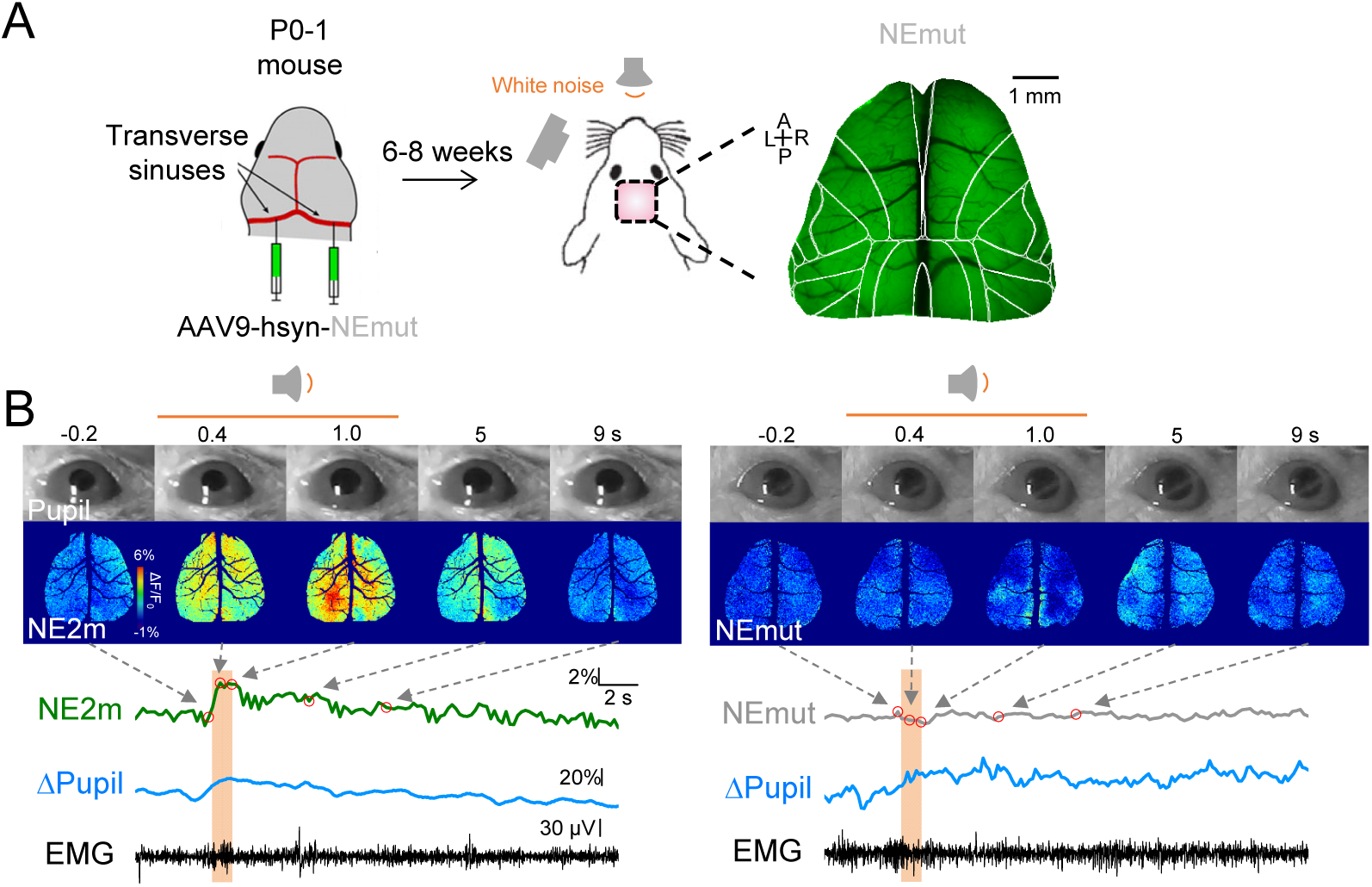
GRAB_NE2m_ and GRAB_NEmut_ fluorescence measure during audio stimulation (related to Figure 5). (A) Schematic diagram depicting the delivery of AAV in P0-P1 mouse pups by injection into the transverse sinuses in P0-P1 mouse for expressing GRAB_NEmut_ in neurons in the dorsal cortex. Also shown are an image of GRAB_NEmut_ fluorescence and the paradigm used for audio stimulation using white noise. (B) Representative images and time course of the change in diameter pupil, GRAB_NE2m_ (left) and GRAB_NEmut_ (right) fluorescence measured in the cortex, and the EMG recording. The shaded areas indicate the delivery of white noise.

**Figure S3.**
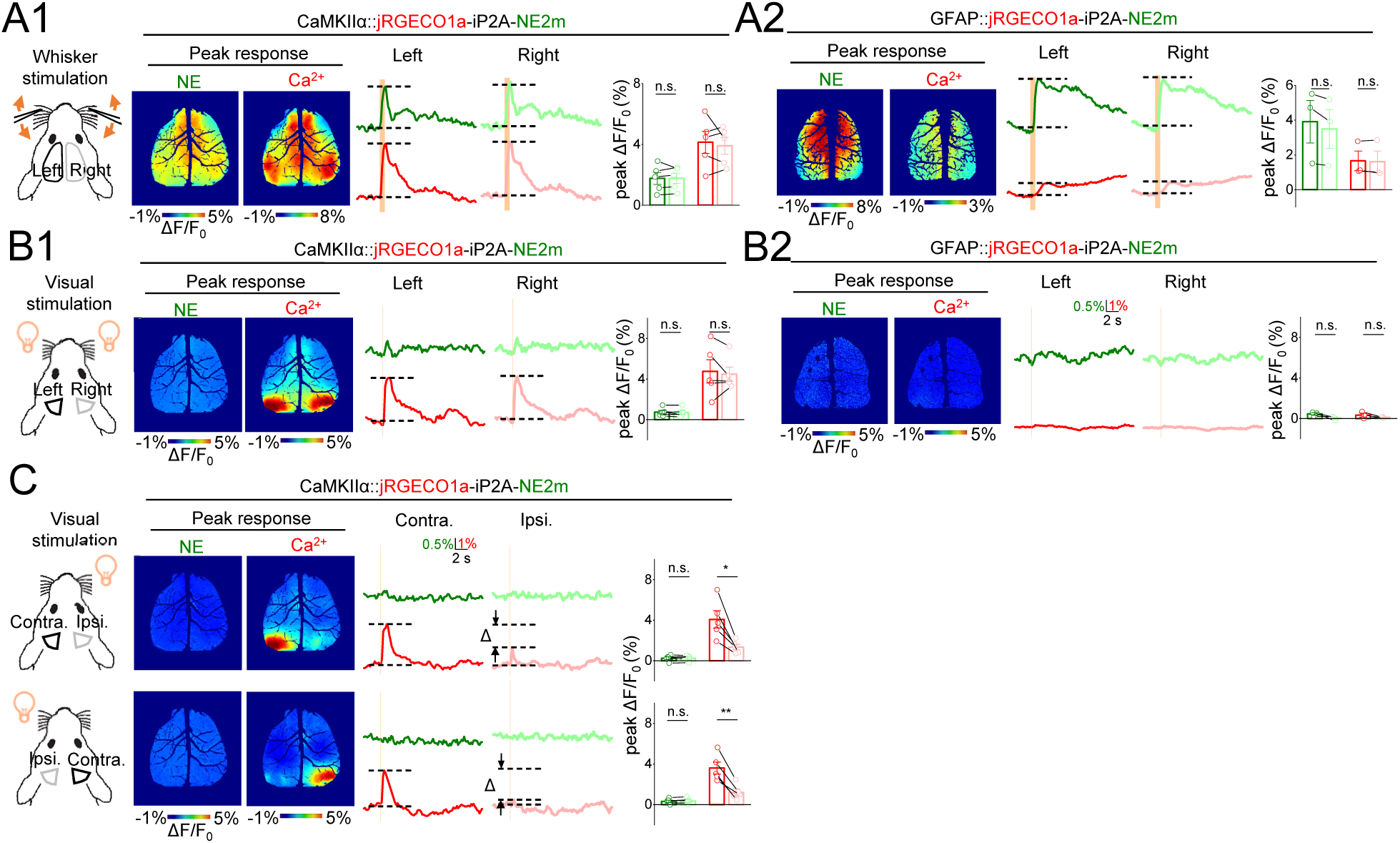
Mesoscopic NE and calcium dynamics in dorsal cortex of awake mice (related to Figure 5). (A-C) Illustrations (left) of whisker stimulation and visual stimulation delivered to CaMKIIα::NECa and GFAP::NECa mice. Shown are the peak response images, representative traces, and summary of the peak responses following bilateral (A-B) or unilateral (C) stimulation of the indicated mice. black and grey lines indicate the ROIs used to analyze the representative traces. n = 3-5 animals per group. \*\**p* < 0.01, \**p* < 0.05, and n.s., not significant (Paired student’s *t*-test).

